# Altered synaptic connectivity and brain function in mice lacking microglial adapter protein Iba1

**DOI:** 10.1101/2021.04.23.441023

**Authors:** Pablo J. Lituma, Evan Woo, Bruce F. O’Hara, Pablo E. Castillo, Nicholas E. S. Sibinga, Sayan Nandi

**Author notes:** Corresponding Author(s)*: Sayan Nandi, Ph.D., Department of Developmental and Molecular Biology, Albert Einstein College of Medicine, Bronx, New York 10461, Nicholas E. S. Sibinga, M.D., Department of Medicine, Department of Developmental and Molecular Biology, Albert Einstein College of Medicine, Bronx, New York 10461, Pablo E. Castillo, M.D., Ph.D., Dominick P. Purpura Department of Neuroscience, Department of Psychiatry and Behavioral Sciences, Albert Einstein College of Medicine, Bronx, New York 10461.

## Abstract

Growing evidence indicates that microglia impact brain function by regulating synaptic pruning and formation, as well as synaptic transmission and plasticity. Iba1 (Ionized Ca^+2^-binding adapter protein 1), encoded by the *Allograft inflammatory factor 1* (*Aif1*) gene, is an actin-interacting protein in microglia. Although Iba1 has long been used as a cellular marker for microglia, its functional role remains unknown. Here, we used global Iba1-deficient (*Aif1*^*-/-*^) mice to characterize microglial activity, synaptic function and behavior. Microglial imaging in acute hippocampal slices and fixed tissues from juvenile mice revealed that *Aif1*^*-/-*^ microglia display reductions in ATP-induced motility and ramification, respectively. Biochemical assays further demonstrated that *Aif1*^*-/-*^ brain tissues exhibit an altered expression of microglial-enriched proteins associated with synaptic pruning. Consistent with these changes, juvenile *Aif1*^*-/-*^ mice displayed deficits in excitatory synapse number and synaptic transmission assessed by neuronal labeling and whole-cell patch-clamp recording in acute hippocampal slices. Unexpectedly, microglial synaptic engulfment capacity was diminished in juvenile *Aif1*^*-/-*^ mice. During early postnatal development when synapse formation is a predominant event in the hippocampus, excitatory synapse number was still reduced in *Aif1*^*-/-*^ mice. Together these findings support an overall role of Iba1 in excitatory synaptic growth in juvenile mice. Lastly, postnatal synaptic deficits persisted in the adulthood and correlated with significant behavioral changes in adult *Aif1*^*-/-*^ mice, which exhibited impairments in object recognition memory and social interaction. These results suggest that Iba1 critically contributes to microglial activity underlying essential neuro-glia developmental processes that may deeply influence behavior.

**Significance:** Abnormal microglia-neuron interaction is increasingly implicated in neurodevelopmental and neuropsychiatric conditions such as autism spectrum disorders and schizophrenia, as well as in neurodegenerative disorders such as Alzheimer’s disease. This study demonstrates that deletion of the microglia-specific protein Iba1, which has long been utilized as a selective microglial marker but whose role has remained unidentified, results in microglial structural and functional impairments that significantly impact synaptic development and behavior. These findings not only highlight the importance of microglia in brain function but may also suggest that modifying microglial function could provide a therapeutic strategy for treatment of neurodevelopmental, neuropsychiatric and neurodegenerative disorders.

## Introduction

Microglia are brain macrophages derived from yolk-sac progenitors, and classically assigned with roles in immune surveillance and response to injury or disease states (1, 2). Recent views, however, indicate that microglia can regulate brain function by remodeling neuronal circuitry both in the healthy and diseases brain (3, 4). For example, excessive complement activity leading to an elevated synaptic pruning is implicated in schizophrenia development (5), and cell culture experiments from schizophrenia patient-derived microglia and neurons exhibit an enhanced microglial pruning of dendritic spines (6). A large multi-level study of transcriptional regulation reveals that a subset of microglial-enriched genes is strongly upregulated in autism spectrum disorders (ASD) peaking during early development, whereas the same module is downregulated in patients with schizophrenia or bipolar disorder after age ∼30 (7). An independent study has further demonstrated alterations in gene expression of specific cell-types including microglia from human patients diagnosed with ASD (8). Thus, understanding the contribution of microglia-neuron interactions to mental health is crucial.

Mice deficient in complement receptor 3 (CR3/CD11b) and fractalkine receptor (CX3CR1) have provided insights into the role of microglia in synaptic pruning of retinal ganglionic cell axon terminals and of dendritic spines in CA1 hippocampus, respectively (9-11). Furthermore, loss of synaptic pruning in CA1 hippocampus correlates with behavioral alterations in sociability and repetitive behavior reminiscent of an ASD phenotype in mice (12, 13). Inappropriate microglial activation and enhanced synaptic pruning are also correlated with cognitive deficits in a mouse model of Down Syndrome (14). Moreover, microglia perform contact-independent synaptic pruning and potentially affect synaptic transmission through neuronal activity-driven release of a TNF-family cytokine, TWEAK by microglial cells (15, 16). On the other hand, microglia can contribute to synapse formation in the developing CA1 hippocampus and somatosensory cortex and in the adult motor cortex, in part by neuronal activity-dependent, as well as by basal release of BDNF and IGF1 (17-19). Another mechanism supporting synapse formation implicated extracellular matrix remodeling by microglia (20). Lastly, neuronal activity triggered by animal experience can recruit microglia for structural and functional modifications of neurons (18, 21-25). Despite these advances, specific contributions of microglia in neurodevelopment and circuit remodeling are not clearly understood.

Iba1 (Ionized Ca^+2^-binding adapter protein 1) encoded by the *Allograft inflammatory factor 1* (*Aif1*) gene, is a conserved intracellular Ca^+2^-binding adapter protein of proinflammatory nature, that is selectively expressed by microglia and macrophages, and has long been utilized as a microglial marker (26-28). *In vitro* overexpression studies have shown that Iba1 interacts with the actin cytoskeleton in membrane ruffles and activates Rac GTPase signaling (29-31). Mice lacking Iba1 (*Aif1*^*-/-*^ mice) appear grossly normal, breed well and show enhanced protection from the development of inflammation (32, 33). In addition, macrophages lacking Iba1 display migration and phagocytosis deficits and secrete reduced levels of proinflammatory cytokines (34). While these *in vitro* studies identify Iba1 functions relevant for microglia, the *in vivo* roles of Iba1 in microglia and in normal brain function are largely unknown.

In this study, we specifically investigated whether microglia lacking Iba1 display any abnormalities and probed for potential alterations in synaptic properties of Iba1-deficient mice. Our findings suggest that Iba1 is required for normal microglial morphology and motility. Importantly, Iba1-deficient mice display strong, persistent synaptic deficits in the CA1 hippocampus, as well as behavioral alterations in object recognition memory and social interaction. Furthermore, our findings support a preferential role for Iba1 in synaptic formation over pruning in juvenile mice. Our study establishes Iba1 as an important modifier of microglia structure and function, and microglia-synapse interactions. To our knowledge, this study is the first report on the role of Iba1 protein in healthy brain.

## Results

### Iba1 contributes to postnatal microglial activity

Previous work showed that *Iba1* mRNA expression is developmentally regulated, with highest expression observed during the first two postnatal weeks followed by a drop in young adult stages (17). In addition, an influx or expansion of microglia in the brain occurs during the second postnatal week, particularly in the hippocampus, coinciding with extensive synaptic remodeling (35-37). Together, these observations are suggestive of important role of microglia during development. Thus, in this study, we primarily focused on microglial development and function between the 2^nd^ and 3^rd^ postnatal weeks.

To begin characterizing microglia in *Aif1*^*-/-*^ mice, we first assessed their presence in P16-P19 (juvenile) *Aif1*^*-/-*^ mice by immunostaining for a microglia-specific protein, P2RY12 (38). We observed that in various areas of *Aif1*^*-/-*^ brains, microglia were present and maintained at comparable densities as in brains of wild type mice (Fig. 1A,B). Next, to determine whether loss of Iba1 altered expression of microglial-enriched proteins, we performed western blotting for CD11b, TREM2 and CX3CR1 (associated with synaptic pruning) and P2RY12 and MafB (associated with maturation and homeostasis) using juvenile whole brain lysates (9-11, 21, 39, 40) (Fig. 1C). We observed that synaptic pruning-associated markers were either upregulated (CX3CR1 and TREM2), or downregulated (CD11b) in the *Aif1*^*-/-*^ brain tissues (Fig. 1C,D). Given that both microglial and peripheral macrophage densities were unchanged in *Aif1*^*-/-*^ mice, the changes in receptor expression likely reflected alterations of protein level per cell in *Aif1*^*-/-*^ mice (Fig. 1) (33). On the other hand, we found no changes in the level of P2RY12 and MafB in *Aif1*^*-/-*^ brains (Fig. 1C,D). These results suggest that while Iba1 is dispensable for microglial generation, it may be required for microglial function.

**Fig. 1.**
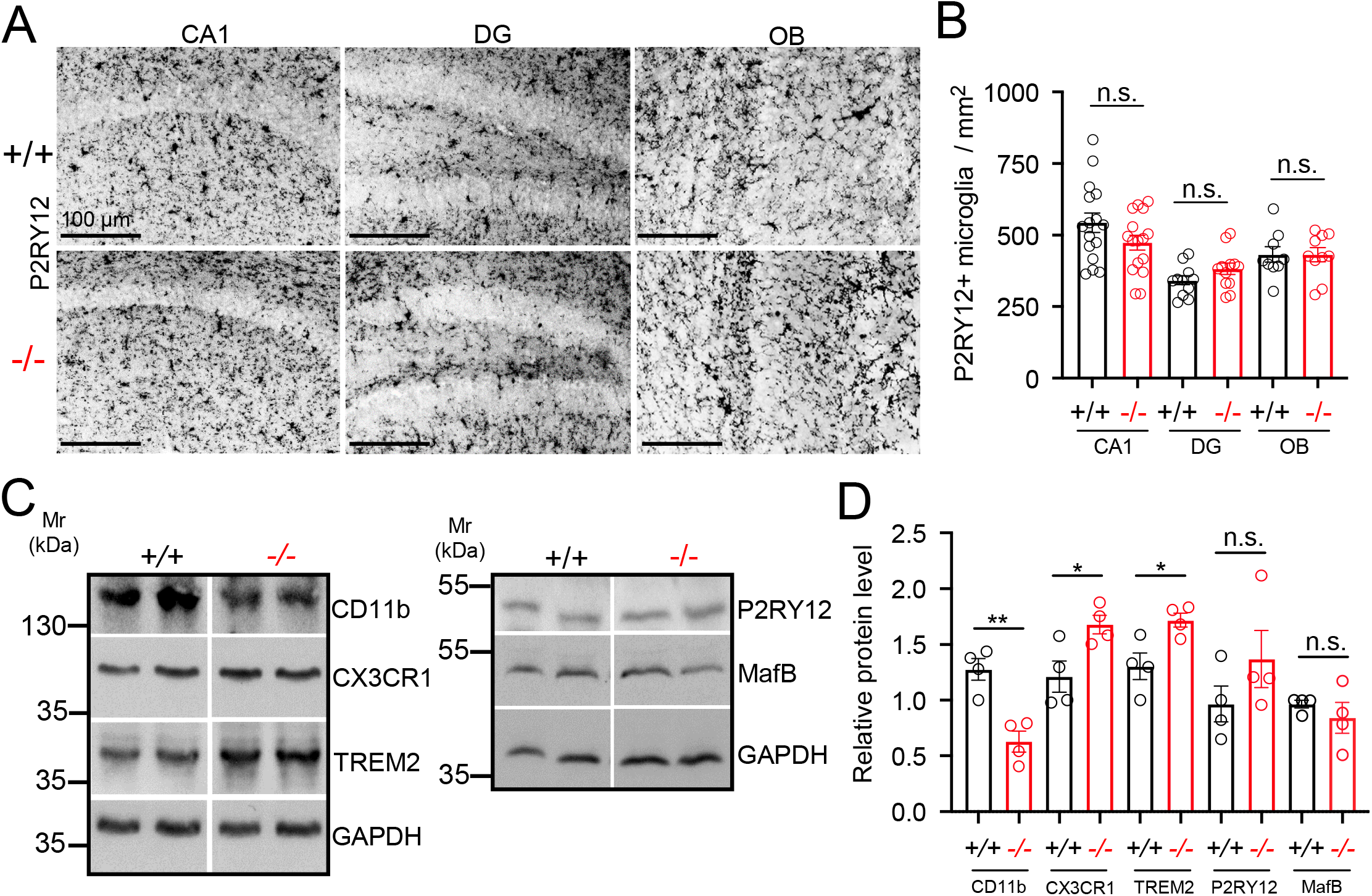
Juvenile *Aif1*^*-/-*^ mice show normal microglial density but an altered whole brain expression of microglial markers of synaptic pruning. *(A)* Photomicrograph of P16-P19 fixed frozen brain sections immunostained with an anti-P2RY12 antibody. DG, dentate gyrus and OB, olfactory bulb. *(B)* Quantitation of P2RY12^+^ microglia in (A). n=10-15 fields per region from 5 mice each genotype, Unpaired Student’s t test. *(C)* Western blotting of P16-P19 whole brain whole cell lysates using antibodies against microglial-enriched synaptic remodeling associated (CD11b, TREM2 and CX3CR1) and homeostatic and maturational (P2RY12 and MafB) proteins. *(D)* Quantitation of western blots in (*C*). n=4 mice per genotype, Unpaired Student’s *t-*test. Average ± SEM. *, p<0.05 and **, p<0.01. n.s.=not significant.

Given that CA1 microglia have previously been implicated in developmental synaptic remodeling (11-13, 19), we focused our attention to this hippocampal area. *Aif1*^*-/-*^ microglia in CA1 were less ramified (but not necessarily more amoeboid) when compared with wild type microglia in morphometric analyses using P2RY12 labeling (Fig. 2A,B). Microglial morphological deficit associated with Iba1 loss was also observed in other brain regions such as neocortex (Fig. 2A,B). Microglial ability to interact with neurons and to modulate synaptic plasticity depends largely on their dynamic process motility (23-25). Previous studies in cultured macrophages and microglial cell lines using an *Aif1* overexpression system suggested that Iba1 localizes with actin cytoskeleton in the membrane ruffles and facilitates motility (29-31). Thus, we investigated whether microglia in acute hippocampal slices display any abnormal motility upon genetic loss of *Aif1*. Using 2-photon microscopy, we acquired z-stack images of CA1 microglia labeled with Isolectin B4 (IB4), before and after focal ATP (1 mM) application using a patch-type pipette as previously described (40-42) (Fig. 2C). We quantified movement of existing processes within a designated region of interest (ROI) around the pipette tip (see SI Materials and Methods) and found that microglia in wild type mice exhibited a greater process motility following ATP application as compared with *Aif1*^*-/-*^ microglia (Fig. 2C,D).

**Fig. 2.**
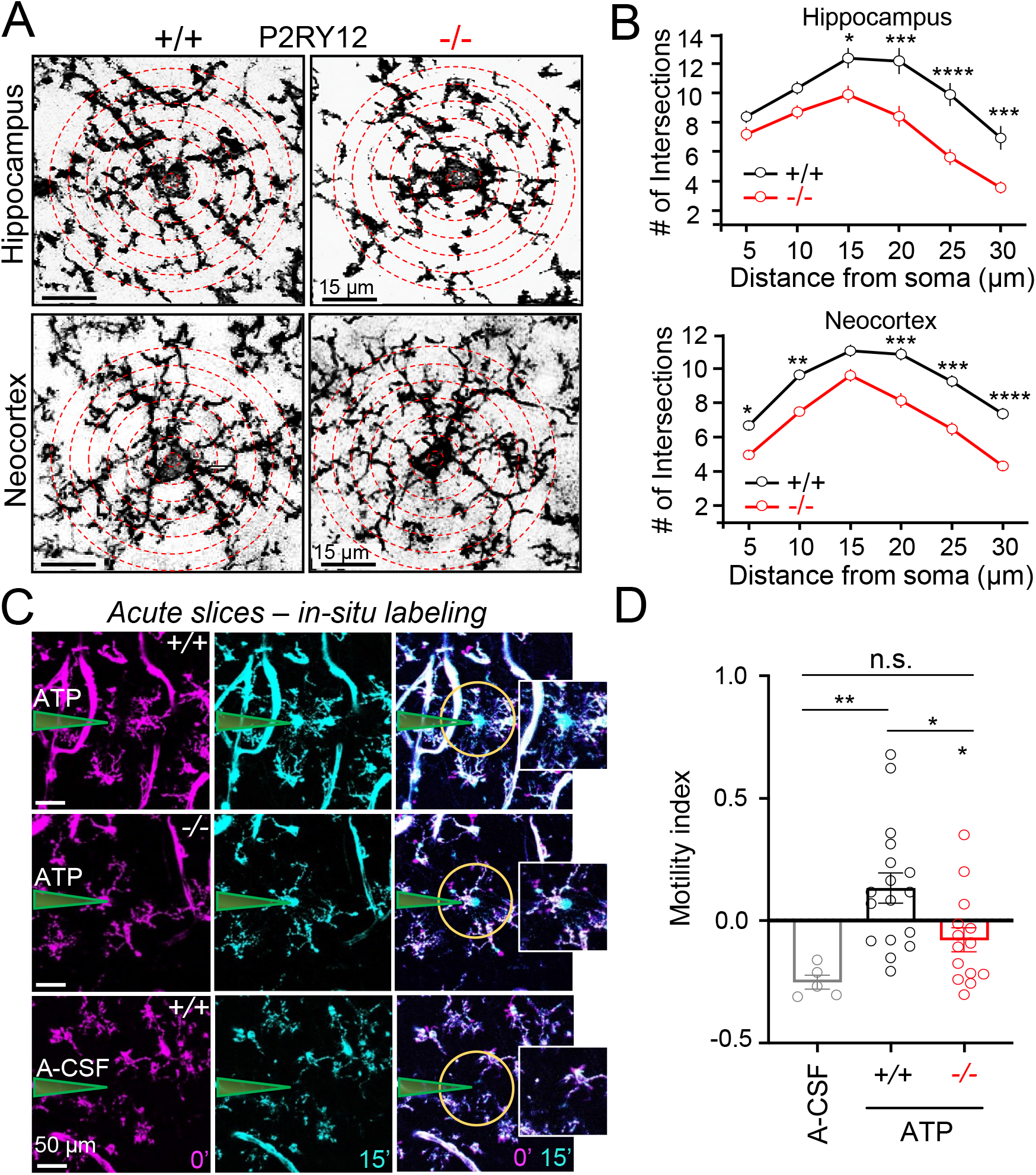
*Aif1*^*-/-*^ microglia in the juvenile brain display reductions in branch complexity and ATP-induced motility. *(A)* Photomicrograph of P16-P19 fixed frozen brain sections immunostained with an anti-P2RY12 antibody facilitated assessment of microglia morphology in hippocampus *(top)* and neocortex *(bottom)*. Sholl analyses was performed by counting number of branch intersections for each of the concentric dotted circles at 5 μm intervals starting from the soma. *(B)* Quantitation of microglia branch complexity revealed significant reductions in hippocampus and neocortex of *Aif1*^*-/-*^ microglia. n=32-45 cells from 8-9 fields per region from 3 mice each genotype; Averages of every intersection were compared between wild type and mutant group in a two-way ANOVA correcting for multiple comparisons using Sidak’s multiple comparison test; hippocampus; F_1,372_=59.58; p<0.0001 and neocortex; F_1,492_=77.15; p<0.0001. *(C)* Representative 2-photon images of microglial motility induced by 15 min of ATP application in wild type *(top)* and *Aif1*^*-/-*^ mice *(middle)*. Microglia were stained with Alexafluor-594-conjugated IB4 in acute hippocampal slices. Note, IB4 also labels blood vessels which have distinct morphological appearances. A-CSF in the pipette served as a negative control *(bottom). (D)* Quantitation of process motility in a demarcated region of interest *(yellow circle)* indicated a significant deficit in *Aif1*^*-/-*^ microglial motility. Motility index (+/+: 0.13 ± 0.06; -/-: −0.08 ± 0.05; p=0.01) was calculated as described in the SI Materials and Methods section. n=18 slices from 6 mice per genotype, Unpaired Student’s t test. Average ± SEM. *, p<0.05, **, p<0.01, ***, p<0.001 and ****, p<0.0001. n.s.=not significant.

These results suggest that Iba1 is required for microglial branching complexity and dynamic process motility, implying that Iba1 may contribute to important microglia-neuron interactions in the juvenile brain.

### Excitatory synapse number and synaptic transmission are reduced in *Aif1*^*-/-*^ mice

Our morphological and biochemical analyses suggested that synaptic remodeling might be affected in juvenile *Aif1*^*-/-*^ mice. To test this possibility, we focused on CA1 pyramidal neurons that have previously been shown to be affected by genetic manipulation in microglia (11-13, 19). We patch-loaded CA1 pyramidal neurons in juvenile mice with biocytin and performed streptavidin-Alexafluor-647-labeling *post-hoc* to assist confocal microscopy (Fig. 3A). Spine density assessed in *stratum radiatum* was significantly reduced in *Aif1*^*-/-*^ mice, suggesting a reduction in excitatory synapse number (Fig. 3B). To test whether this structural change was accompanied by a reduction in excitatory drive, we first monitored miniature excitatory postsynaptic currents (mEPSCs) by performing whole-cell patch-clamp of CA1 pyramidal neurons in the presence of tetrodotoxin (0.5 µM) and the GABA_A_ receptor antagonist, picrotoxin (100 µM) (Fig. 3C). Synaptic events were registered and both current amplitude and frequency was assessed in wild type and *Aif1*^*-/-*^ mice (Fig. 3C,D). While amplitude of mEPSC was unaltered, frequency was significantly diminished in *Aif1*^*-/-*^ mice when compared with wild type mice (Fig. 3D). To determine changes in evoked excitatory transmission, we performed extracellular field recordings of Schaffer collateral-CA1 synapses and found reduced synaptic strength in *Aif1*^*-/-*^ mice as compared to control, as indicated by a reduction of Input/output function (Fig. S1A,B). In addition, paired-pulse ratio at varying stimulus intervals showed no significant changes, suggesting normal presynaptic function in *Aif1*^*-/-*^ mice (Fig. S1C,D).

**Fig. 3.**
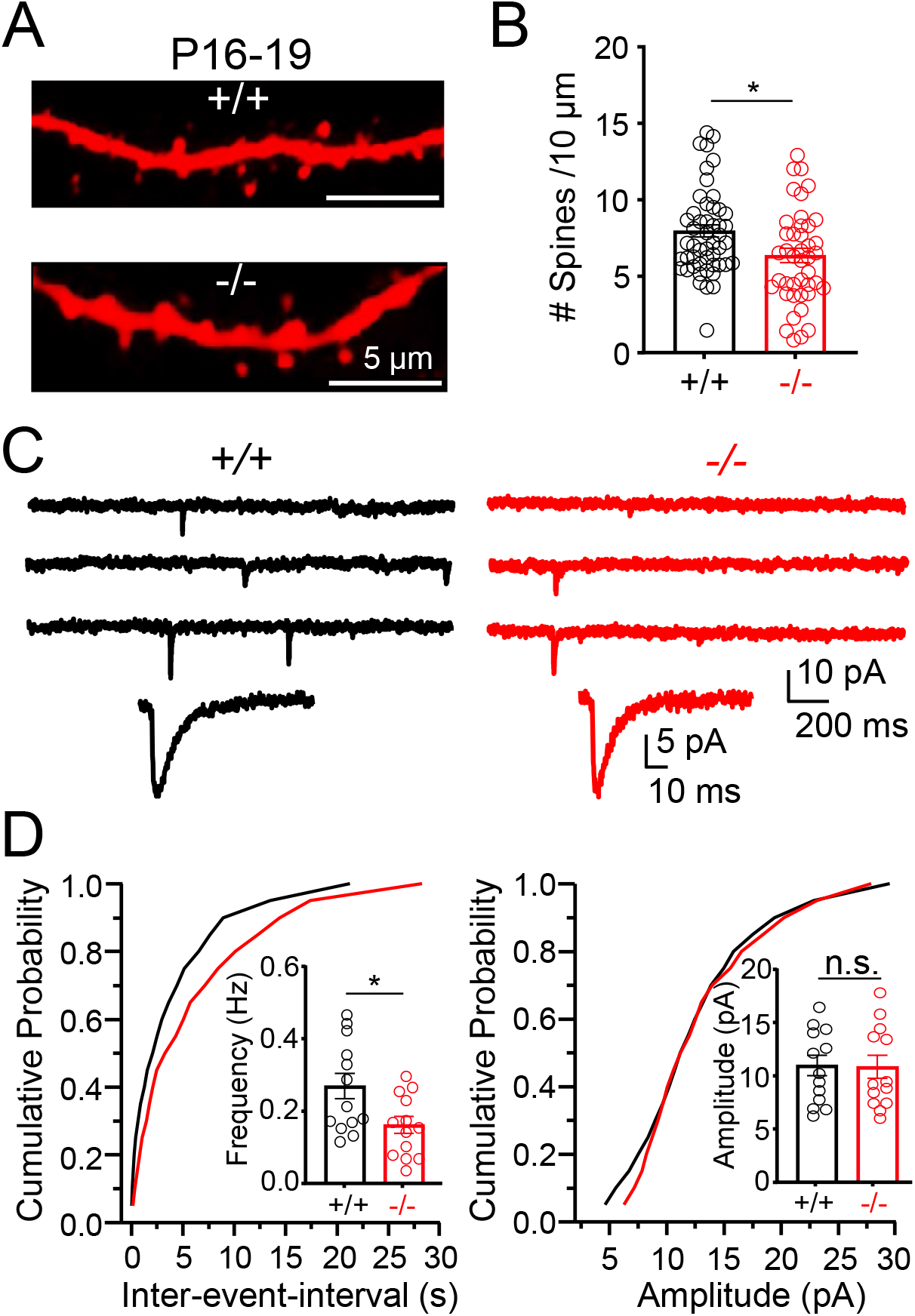
Reduction in excitatory synapses in juvenile *Aif1*^*-/-*^ mice. *(A)* CA1 pyramidal neurons in P16-P19 mice were patch-loaded with biocytin, fixed and stained with Alexafluor-647-conjugated streptavidin. Spine images were acquired from apical dendrites within 100 μm distance from soma. *(B)* Quantitation of spine density (Number per 10 µm, +*/*+: 7.93 ± 0.39; *-/-*: 6.47 ± 0.47; p=0.02). n= 58 (+*/*+) and 46 (*-/-*) dendrites from 13 cells from 8 mice per genotype, Unpaired Student’s t test. *(C)* Representative traces of whole-cell patch clamp recording from CA1 pyramidal cells in acute hippocampal slices treated with tetrodotoxin (0.5 µM) and picrotoxin (100 µM). *(D)* Quantitation of frequency and amplitude of mEPSC (frequency Hz, +*/*+: 0.27 ± 0.03; *-/-*: 0.16 ± 0.02; p=0.02). n=13 cells from 6 mice per genotype, Unpaired Student’s t test. Average ± SEM. *, p<0.05. n.s.=not significant.

To assess whether microglial deficits persist in the adulthood, we performed microglial Sholl and immunoblotting analyses for microglial morphology and synaptic remodeling-associated proteins, respectively, in 3-month-old wild type and *Aif1*^*-/-*^ mice. Expression of CD11b, TREM2 and CX3CR1 proteins was normal in 3-month-old *Aif1*^*-/-*^ brain tissues (Fig. S2A,B). In fact, CA1 microglial morphology in adult *Aif1*^*-/-*^ mice was more complex compared to wild type microglia (Fig. S2C,D), suggesting that compensatory changes have occurred post-development to normalize some aspects of microglial activity in *Aif1*^*-/-*^ brains. We further assessed synaptic structure-function as described above (Fig. 3), but in 3-month-old wild type and *Aif1*^*-/-*^ mice (Fig. 4). Intriguingly, we observed deficits in both synaptic structure (Fig. 4A,B) and function (Fig. 4C,D) in adult *Aif1*^*-/-*^ mice when compared with wild type mice.

**Fig. 4.**
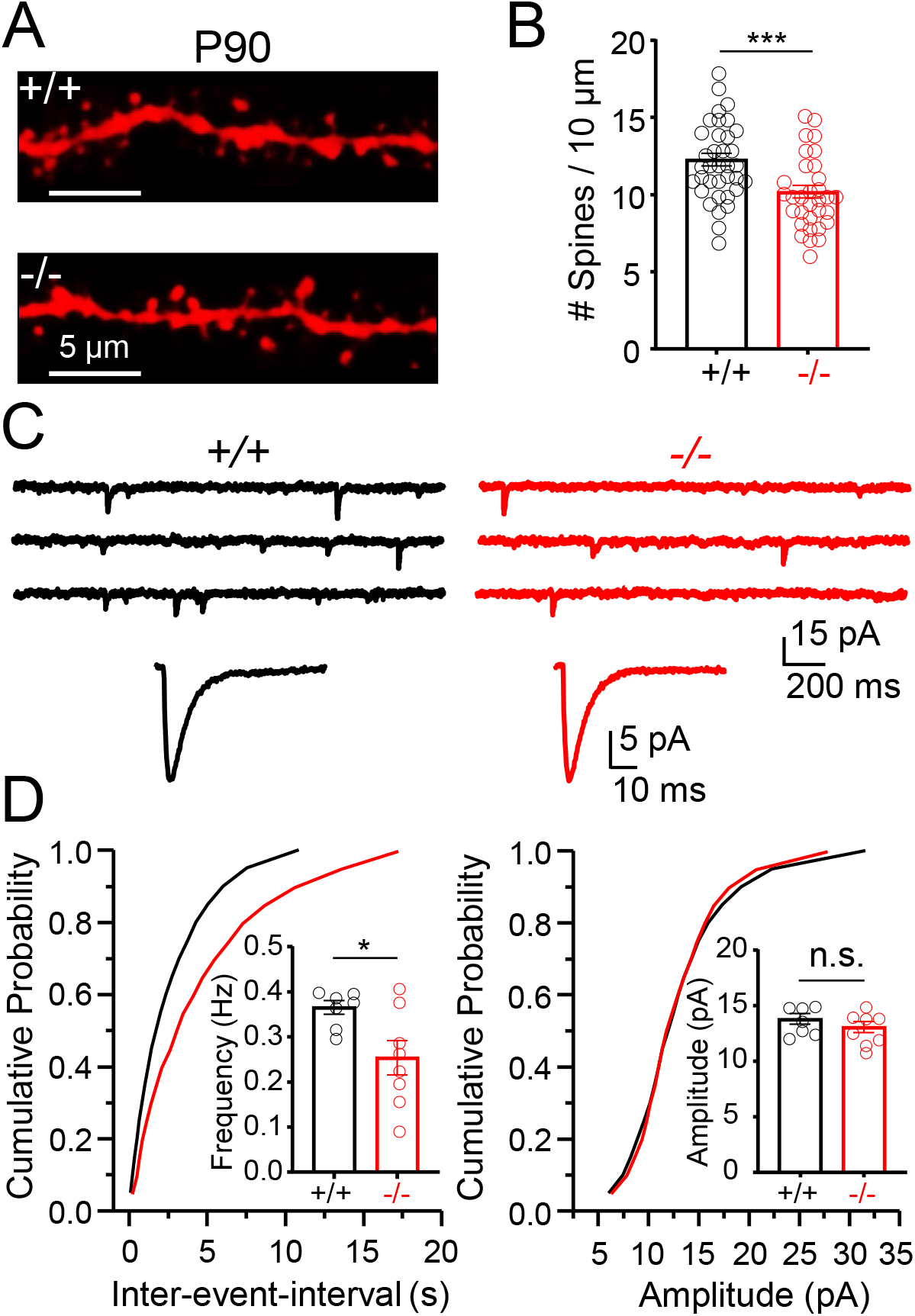
Deficit in excitatory synapse density persists in adult *Aif1*^*-/-*^ mice. *(A)* CA1 pyramidal neurons in P85-90 mice were patch-loaded with biocytin, fixed and stained with Alexafluor-647-conjugated streptavidin. Spine images were acquired from apical dendrites within 100 μm distance from soma. *(B)* Quantitation of spine density (Number per 10 μm, +*/*+: 12.25 ± 0.41; *-/-*: 10.18 ± 41; p=0.0007). n=36 (+*/*+) and 32 (*-/-*) dendrites from 7-8 cells from 4 mice per genotype; Unpaired Student’s *t*-test. *(C)* Representative traces of whole-cell patch clamp recording from CA1 pyramidal cells in acute hippocampal slices treated with tetrodotoxin (0.5 μM) and picrotoxin (100 μM). *(D)* Quantitation of frequency and amplitude of mEPSC (frequency Hz, +*/*+: 0.37 ± 0.02; *-/-*: 0.25 ± 0.04; p=0.02). n=7-8 cells from 4 mice per genotype, Unpaired Student’s t test. Average ± SEM. *, p<0.05 and ***, p<0.001. n.s.=not significant.

Taken together, our evaluation of excitatory transmission revealed that synaptic function was significantly altered in juvenile *Aif1*^*-/-*^ mice, at least in part as a result of synaptic structural deficits observed in these mice. In addition, synaptic structure-function deficits persisted in adult *Aif1*^*-/-*^ mice despite microglial deficits were largely compensated post-development.

### Adult *Aif1*^*-/-*^ mice display behavioral alterations

Microglial dysfunction has been correlated with alterations in mnemonic processes and social behavior in mice (12, 13, 43). Based on our synaptic physiology analyses involving CA1 hippocampus, we anticipated that relevant behavior might be affected in adult *Aif1*^*-/-*^ mice. Since *Aif1*^*-/-*^ mice have not been formally characterized in behavioral terms, we first assessed the general locomotion/exploration and anxiety using the open field test (44, 45) (Fig. S3A). Compared with wild type mice, *Aif1*^*-/-*^ mice traveled a greater distance (by 19%) in the open field, suggesting a modest increase in exploration/locomotion in the latter (Fig. S3A). We further subjected *Aif1*^*-/-*^ mice to the elevated plus maze test, as a definitive measure of anxiety-like behavior (46, 47) (Fig. S3B). *Aif1*^*-/-*^ and wild type mice spent an equivalent amount of time in the open arm, indicating no differences in anxiety (Fig. S3B). Furthermore, *Aif1*^*-/-*^ mice did not preferentially avoid the center area during the open field test, consistent with an absence of anxiety-like behavior in these mice (Fig. S3A). Together, these studies show that *Aif1*^*-/-*^ mice display an increase in general exploration/locomotion but lack anxiety-like behavior.

We next performed the novel object recognition test, a task dependent in part on CA1 hippocampus (48-50). In this test, a mouse was first exposed to two identical objects in an open field, and then after an hour of retention, re-exposed to the open field with one of the objects switched to a new object (51) (Fig. 5A). Wild type mice displayed preference for the novel object over the familiar one upon re-exposure as assessed by the discrimination index whereas such preference was reduced (by 64%) in *Aif1*^*-/-*^ mice (Fig. 5A). Total overall exploration for novel and familiar object combined remained comparable between genotypes (Fig. 5A). These results suggest that adult *Aif1*^*-/-*^ mice display a strong reduction in object recognition memory, consistent with a persistent deficit in hippocampal synaptic connectivity.

**Fig. 5.**
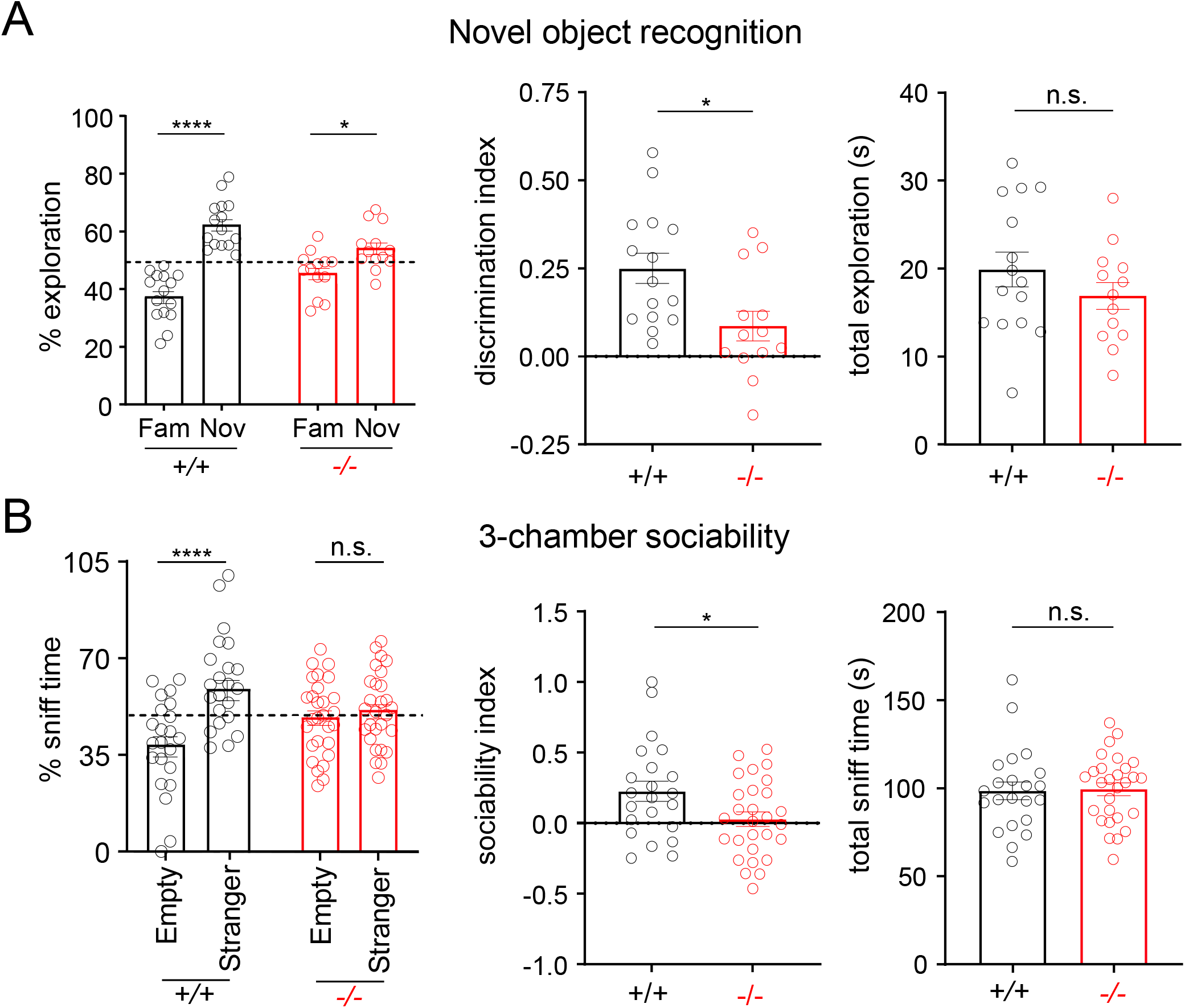
Adult *Aif1*^*-/-*^ mice display behavioral alterations. *(A)* Mice explored an open field guided by visual cues; first in the presence of two identical objects (3 min of training) and an hour later in the presence of two objects when one of the two old objects was replaced with a new object (3 min of testing). n=15 (+*/*+) and 13 (*-/-*); Percentage exploration for each object during testing was calculated and statistical significance across different conditions and genotypes was assessed by a two-way ANOVA correcting for multiple comparisons using Tukey’s multiple comparison test *(left)*. Discrimination index during testing was calculated (+/+: 0.25 ± 0.04; -/-: 0.09 ± 0.04; p=0.01) as described in SI Materials and Methods section *(middle)*. Total exploration during testing (+/+: 19.88 ± 1.96 s; -/-: 16.89 ± 1.54 s; p=0.23) was recorded *(right)*. Statistical significance between two genotypes was assessed using unpaired Student’s t test. *(B)* Mice explored a 3-chamber plexiglass with sliding doors; first alone in center chamber (5 min) with doors closed, then with a stranger mouse of same sex which was placed in a circular cage in one of the two chambers (10 min) with both doors open (for sociability). n=23 (+*/*+) and 28 (*-/-*); Percentage sniff time was calculated and statistical significance across different conditions and genotypes was assessed by a two-way ANOVA correcting for multiple comparisons using Tukey’s multiple comparison test *(left)*. Sociability index was calculated (+/+: 0.23 ± 0.07; -/-: 0.03 ± 0.05; p=0.03) as described in SI Materials and Methods section *(middle)*. Total sniff time during each trial (+/+: 98.45 ± 5.1 s; -/-: 99.45 ± 3.62 s; p=0.87) was recorded *(right)*. Statistical significance between two genotypes was assessed using unpaired Student’s t test. Mice were between 3-5 months. Average ± SEM. *, p<0.05 and ****, p<0.0001. n.s.=not significant.

To further investigate potential alterations in neural circuits outside the hippocampus such as cortical or midbrain structures in *Aif1*^*-/-*^ mice, we performed a three-chamber social test (52-54). A test mouse was first assessed for social interaction by exposing it to a stranger mouse, and then for social novelty by additionally exposing it to a second stranger mouse. Since *Aif1*^*-/-*^ mice had an increase in exploration/locomotion in the open field test (Fig. S3A), we assessed sniff time rather than time spent in each chamber as a measure of interaction. During the first trial, wild type mice preferentially sniffed the stranger mice more than the empty chamber, while *Aif1*^*-/-*^ mice sniffed both the stranger mice and the empty chamber equally (Fig. 5B). The sociability index was reduced by 87% in *Aif1*^*-/-*^ mice compared with wild type mice (Fig. 5B). Both groups displayed equivalent total sniff time (Fig. 5B). These results suggest that *Aif1*^*-/-*^ mice exhibit a strong deficit in social interaction and neural circuit impairment may occur in additional *Aif1*^*-/-*^ brain regions.

In summary, adult *Aif1*^*-/-*^ mice showed significant alterations in behavior that are often associated with neurodevelopmental disorders involving multiple brain areas.

### Iba1 facilitates growth of excitatory synapses in the juvenile brain

The effect of Iba1 loss on synaptic remodeling function supported the idea that Iba1 promotes growth of excitatory postsynaptic structures (Figs. 3, 4 and S1). We thus investigated whether the synaptic growth-promoting function of Iba1 could be attributed to an ability to limit synaptic pruning or to facilitate synaptic formation. To assess a role for Iba1 in synaptic pruning, we performed a synaptic engulfment assay using immunostaining and quantitative Airyscan microscopy using P16-P19 fixed frozen tissue sections from wild type and *Aif1*^*-/-*^ mice (12, 21, 55). P2RY12, CD68 and PSD-95 served as respective markers for microglia, microglial lysosome, and excitatory postsynaptic structures. Airyscan microscopy followed by 3D-surface rendering using Imaris software revealed no significant change in the volume of CD68+ structures per *Aif1*^*-/-*^ microglial volume (Fig. 6A,B). Using the “ mask function” of Imaris (see SI Materials and Methods), a 45% reduction in the volume of PSD-95+ structures that were associated with microglial CD68+ structures was observed in *Aif1*^*-/-*^ mice as compared to wild type mice (Fig. 6A,B). Consistent with earlier observations (Fig. 3A,B), we detected a 50% reduction in the volume of overall PSD-95+ structures in hippocampal CA1 fields of *Aif1*^*-/-*^ mice compared with wild type mice (Fig. 6A,B). These results suggest that the reduction in the number of excitatory synapses upon Iba1 loss is not due to an enhanced contact-dependent synaptic pruning. While an increase in microglia-driven contact-independent pruning which requires sensory experience (15, 16), could potentially contribute to the reduction in number of excitatory synapses in *Aif1*^*-/-*^ mice, it is an unlikely scenario given that our synaptic structure-function analyses were performed under basal conditions (Figs. 3, 4 and 6).

**Fig. 6.**
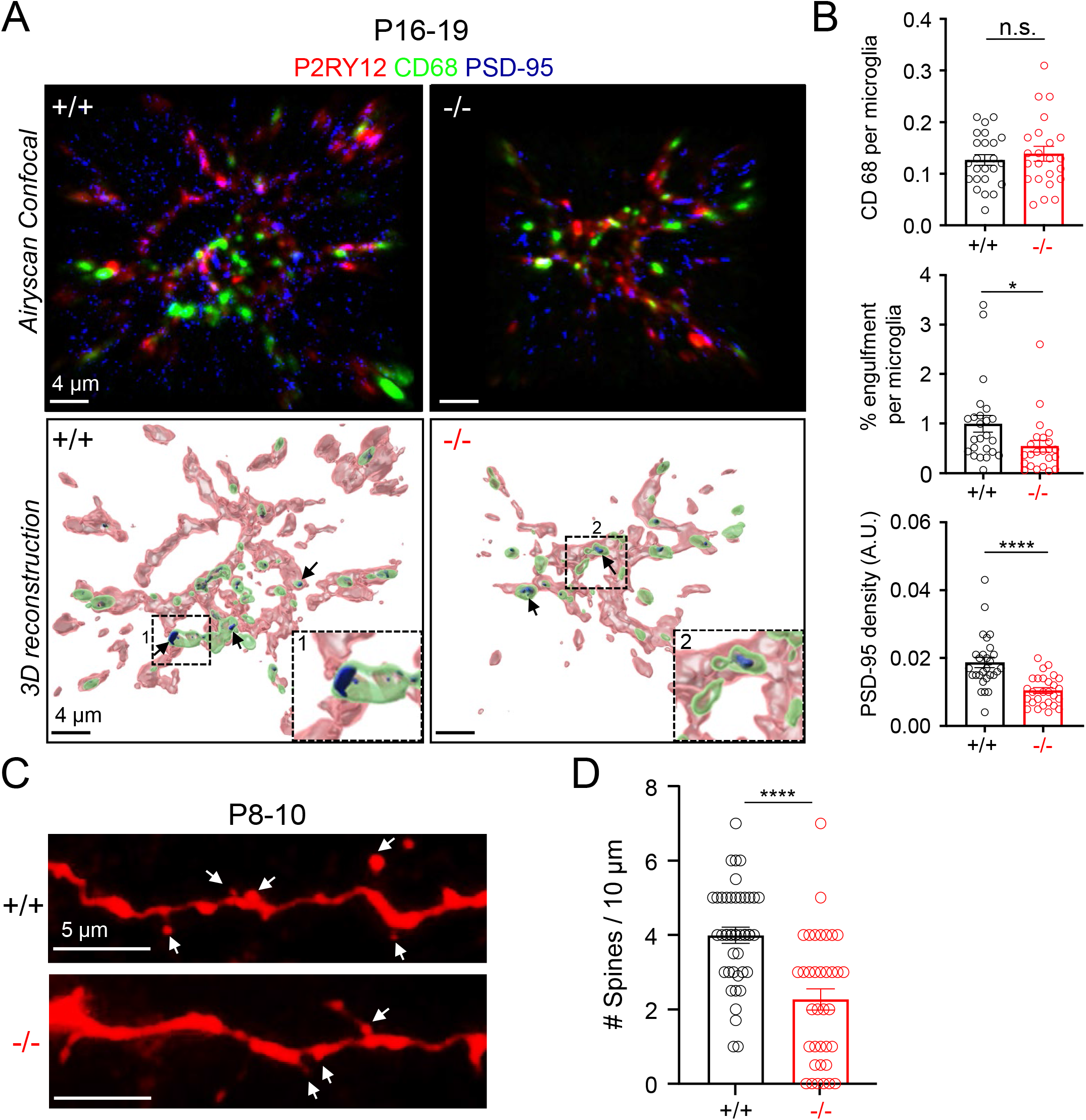
Iba1 loss attenuates microglial synapse engulfment in the juvenile mice but also causes an early synapse formation deficit. *(A)* Confocal z-stack images of CA1 microglia and excitatory synapses in fixed brain tissues from P16-19 mice *(top)*. 3D-surface rendering and using “ mask” function in Imaris demonstrating postsynaptic (PSD-95, blue, arrows) in microglial (P2RY12, red) lysosomal structure (CD68, green) *(bottom)*. Arrows indicate examples of engulfed synaptic materials. *(B)* A similar volume of CD68+ structures per microglial volume (+/+: 0.12 ± 0.01; *-/-*: 0.14 ± 0.01; p=0.5) in wild type and *Aif1*^*-/-*^ mice *(top)*. A reduced percentage volume of engulfed synaptic material (PSD-95+ structures within CD68+ structures) per microglia volume (+/+: 0.99 ± 0.17; *-/-*: 0.54 ± 0.12; p=0.03) in *Aif1*^*-/-*^ mice *(middle)*. n=24 cells from 6 mice per genotype. A reduction in total volume of PSD-95+ structures per CA1 field (+/+: 0.02 ± 0.001; *-/-*: 0.01± 0.001; p<0.0001) in *Aif1*^*-/-*^ mice *(bottom)*. n=28 fields from 4 mice per genotype. *(C)* CA1 pyramidal neurons in P8-10 mice were patch-loaded with biocytin, fixed and stained with Alexafluor-594-conjugated streptavidin. Confocal spine images (arrows) were acquired from apical dendrites within 50 μm distance from soma. *(D)* Quantitation of spine density in P8-10 mice (Number per 10 µm, +*/*+: 4.00 ± 0.22; *-/-*: 2.66 ± 0.29; p<0.0001). n= 41 (+*/*+) and 36 (*-/-*) dendrites from 10 cells from 7 mice per genotype, Unpaired Student’s t test; Average ± SEM. *, p<0.05 and ****, p<0.0001. n.s.=not significant.

Recent studies show that microglia can contribute to excitatory synapse formation in juvenile mice (18, 19). Iba1 could promote synapse formation of cortical neurons during early postnatal development (P8-10) (17). To determine if Iba1 may perform a similar function in CA1 pyramidal neurons, we further assessed dendritic spine density in P8-10 mice, at the time when synaptic pruning events are largely absent (11, 19) (Fig. 6C). Our results revealed a 43% reduction in spine density in P8-10 *Aif1*^*-/-*^ mice compared with wild type mice (Fig. 6D). This result suggests that Iba1 deletion impairs synapse formation.

Lastly, astrocytes can contribute to synaptic formation, pruning, transmission, plasticity, and recent studies position astrocytes downstream of microglia (56-60). We therefore performed density and morphological analyses of CA1 astrocytes using GFAP immunostaining. Assessment of CA1 astrocytic number and morphology failed to reveal significant differences between wild type and *Aif1*^*-/-*^ mice (Fig. S4). While it is plausible that astrocytes could still affect synaptic processes despite their normal number and morphology in the absence of Iba1, this possibility remains to be fully investigated.

Together our results suggest that Iba1 contributes to an overall growth of excitatory synapses, by virtue of its predominant role in synapse formation over pruning in the juvenile brain.

## Discussion

We report that mice lacking Iba1 have altered microglial structure-function, deficits in excitatory synapses, and changes in behavior. Our findings highlight a pivotal contribution of microglial activity during neurodevelopment to synaptic refinement that may influence behavior. To the best of our knowledge, this study provides the first characterization of Iba1 contributions to microglial biology and brain function using a genetic loss-of-function approach.

Microglia are a selective cell population in brain tissue that can migrate or dynamically extend or retract processes either to probe local environments (surveillance) or to respond to pathogens or injury (chemotaxis) (41, 61). In mice with genetic inactivation of Iba1 expression, we identified a significant impairment in microglial ramification and ATP-induced motility in acute brain slices, suggesting that Iba1 may indeed play essential roles in dynamic motility of these cells (Fig. 2). Microglial motility and associated neuronal plasticity are altered by ATP-driven activation of microglial purinergic receptor P2RY12 (23, 25, 41, 42). However, we did not find changes in P2RY12 protein level in *Aif1*^*-/-*^ brain tissue, suggesting that P2RY12 may act upstream of Iba1 to enable motility (Figs. 1 and 2). In addition, focal laser injury can enhance Ca^+2^ activity in microglial soma and protruding processes (62). Although its role in intracellular Ca^+2^ dynamics has not been investigated, Iba1 protein is known to bind Ca^+2^, and to activate Rac GTPase signaling (31, 63) — these Iba1 functions could potentially contribute to microglial process extensions that may deeply influence neuro-glia interactions.

Recent studies demonstrate that microglia are not only essential for synaptic pruning but may also participate in synapse formation during early neurodevelopmental stages (17-19). In the somatosensory cortex, a high *Iba1* mRNA levels have been correlated with synapse formation in early postnatal mice (17). Consistent with this observation, our neuronal structural analysis revealed strong reductions of excitatory synaptic density in the CA1 hippocampus of early postnatal as well as juvenile *Aif1*^*-/-*^ mice (Figs. 3 and 6). Paradoxically, we also observed a reduction in microglial synaptic engulfment capacity in hippocampal CA1 when Iba1 is absent (Fig. 6A,B). Given that synaptic formation precedes pruning and pruning peaks between 2^nd^-3^rd^ postnatal week with concomitant synaptic formation, it is conceivable that of the two remodeling processes, synaptic formation is a more dominant process in the juvenile mice (11, 12, 18, 64). Consistent with a role of Iba1 in synapse formation, *Aif1*^*-/-*^ synaptic deficits were apparent by P8-P10 when synaptogenesis reaches its peak in somatosensory cortex and to a large extent, in hippocampus (17, 65). It is also possible that reduction in the capacity of microglial synaptic engulfment in juvenile *Aif1*^*-/-*^ mice could be in part due to compensatory changes to an early synapse formation deficit in these mice. Future approaches should consider temporal ablation of Iba1 function to better study the role of microglia in defining the “ critical-period plasticity” (36, 37).

How exactly Iba1 regulates microglial function warrants further investigation and a deeper mechanistic understanding of the cellular role of Iba1. Although generally described as a cytosolic protein, Iba1 subcellular localization and function remains to be determined. Iba1 activities within cells may include, but are not limited to 1) actin-based cytoskeletal roles that drive microglial phagocytotic and/or secretory functions such as vesicular exocytosis or exosomal release of cytokines and miRNAs (66), and 2) regulation of microglial expression of genes encoding critical receptors, cytokines and matrix metalloproteases (21, 67). As noted above, we found decreased ramification and motility of microglia lacking Iba1, consistent with altered cytoskeletal functions (Fig. 2). We also identified clear differences in protein levels associated with synaptic pruning in *Aif1*^*-/-*^ brain tissues (Fig. 1), but our observations so far cannot distinguish between possible direct or indirect mechanisms by which Iba1 could regulate specific microglial gene expression during neurodevelopment.

Human studies demonstrate that enhanced Iba1 expression is correlated with the onset of ASD and Down syndrome during early life (7, 14). In addition, Iba1 is downregulated with the onset of schizophrenia and bipolar disorder around age 30 (7). These results are consistent with microglial roles in neurodevelopment and neurophysiology and the idea that their dysfunction correlates with onset of neurodevelopmental and neuropsychiatric disorders. Human epidemiological studies have clearly demonstrated a link between early immune activation and psychiatric conditions in later life including depression and psychosis (68, 69). Furthermore, prenatal infections have been linked to the development of ASD and schizophrenia (70, 71). Our Iba1 loss-of-function model of microglial dysfunction supports the impact of early developmental changes to the alterations of cognitive and social behaviors in adulthood (Fig. 4). These results reinforce the need to deeply assess microglial properties during those early immune perturbations leading up to the development of cognitive dysfunctions later in life. Recent work has demonstrated that combined auditory and visual stimulation to induce gamma oscillations can activate microglia and lead to reductions in amyloid plaques that are associated with cognitive improvements (72). Furthermore, microglial pruning of engram cell synapses correlates with forgetting mechanism in adult mice (43). As microglia function becomes further implicated in neurodevelopmental, neuropsychiatric and neurodegenerative disorders, a better understanding of neuro-glia interactions will provide new therapeutic insights.

## Materials and Methods

*Aif1*^*-/-*^ mice were generated, maintained and bred as previously described (32, 33) and were further backcrossed to C57BL/6J for more than 10 generations (34). All experimental procedures involving mice were carried out following the guidelines of IACUC at the Albert Einstein College of Medicine. Information on antibodies and procedures involving IHC, hippocampal slice preparation, electrophysiology, behaviour, microglial morphology and motility, biochemical evaluation of brain tissues were detailed in SI Material and Methods. In all cases, the investigators were blind to the conditions and genotypes during data acquisition and analysis.

## Acknowledgments

We thank Dr. David Julius, UCSF for P2RY12 antibody. We thank Dr. Kostantin Dobrenis, Kevin Fisher and Vladimir Mudragel of Einstein Neural Cell Engineering and Imaging Core supported by Rose F. Kennedy Intellectual and Developmental Disabilities Research Center (IDDRC) for technical advice and acknowledge support from NIH shared instrument grant S10OD025295 K.D. We thank Dr. Derek M. Huffman and Zunju Hu of Healthy Aging Physiology Core, Einstein and Deann Hopkins of Rodent Behavior Core, University of Kentucky for assistance with behavioral analyses, Dr. Prameladevi Chinnasamy and Smitha Jayakumar for assistance with mouse breeding, Kayla Oriyomi and Grace Tremonti for assistance with sholl analyses, and the members of the Castillo laboratory for providing critique of this work. This work was supported by NIH grants R21 NS116480 and R21 MH124294 and Einstein Nathan Shock Center P&F award to S.N., R01 HL128066 and R01 HL133861 to N.E.S., R01 MH125772, R01 MH116673, and R01 NS113600 to P.E.C. and a Ruth L. Kirschstein NRSA Fellowship F31MH109267 to P.J.L.

## Conflict of interest

Authors declare no conflict of interest.

## Author contributions

S.N. and P.J.L. designed studies, performed experiments, analyzed data and wrote the manuscript; S.N., N.E.S.S., P.E.C. and B.F.O supervised experiments; N.E.S.S. and P.E.C. assisted in manuscript preparation; E.W. performed experiments.

## Supplementary Information for

## SI Materials and Methods

### Antibodies

The following antibodies were used for IHC: P2RY12 (1:200, rabbit polyclonal, AnaSpec) (1), CD68 (1:200, rat monoclonal IgG_2a_, Bio-Rad) (2), PSD-95 (1:100, mouse monoclonal IgG_2aκ,_ Sigma-Millipore) (3) and Iba1 (1:700, rabbit polyclonal, Wako) (4). The following antibodies were used for immunoblotting: TREM2 (1:1000; rabbit monoclonal; Cell Signaling) (5), CX3CR1 (1:100; mouse monoclonal IgG_2a_; Santa Cruz Biotechnology), CD11b (1:1000, rabbit monoclonal, Abcam) (6), MafB (1:100, mouse monoclonal IgG_2b_, Santa Cruz Biotechnology) (7), P2RY12 (1:1000, rabbit polyclonal), a gift from Dr. David Julius, University of California San Francisco (8), Iba1 (1:2000, rabbit monoclonal, Abcam) (9) and GAPDH (1:10000, rabbit polyclonal; Santa Cruz Biotechnology).

### Immunohistochemistry and Microscopy

For fixed-frozen brain sections, P16-P19 wild type and *Aif1*^*-/-*^ mice were transcardially perfused first by pre-chilled heparinized PBS 1X followed by 4% paraformaldehyde (PFA) in PBS 1X (pH 7.4) (10). Brains were removed from skulls, cut mid-sagittally and incubated further with 4% PFA in PBS 1X overnight at 4° C. Brains were subsequently incubated first with 10% sucrose in PBS 1X for 6 h at 4° C and then with 20% sucrose in PBS 1X for overnight at 4° C. Brains were embedded using TissuePlus O.C.T. compound (Fisher Scientific, MA) and stored at −80° C. Fixed-frozen brains were cut to generate 14-μm or 30-μm thick sections using a cryostat (Leica CM 1900). Sections were incubated with primary antibodies overnight at 4° C and then with Alexafluor-conjugated secondary antibodies (ThermoFisher Scientific, MA) for 1.5 h at 4° C before mounting using ProLong Diamond Antifade Mountant with DAPI (ThermoFisher Scientific, MA). For fixed-floating brain sections, adult wild type and *Aif1*^*-/-*^ mice were sacrificed, and acute 300-μm thick sections were obtained using a vibratome. Sections were incubated further with 4% PFA in PBS 1X overnight at 4° C. Fixed-floating sections were incubated with primary antibodies overnight at 4° C and then with Alexafluor-conjugated secondary antibodies (ThermoFisher Scientific, MA) overnight at 4° C in a 24-well plate (Corning Costar) before mounting using ProLong Diamond Antifade Mountant with DAPI (ThermoFisher Scientific, MA).

### Tissue homogenate preparation

Brains from P16-P19 wild type and *Aif1*^*-/-*^ mice were dissected mid-sagittally and halves were homogenized on ice using 1 ml of pre-chilled homogenization buffer containing 50 mM Tris, 150 mM NaCl, 1mM EDTA and 1% NP-40, pH 7.4 in the presence of reconstituted protease inhibitor cocktail (MilliporeSigma, MA) and PhosSTOP phosphatase inhibitor cocktail (MilliporeSigma, MA) in Dounce homogenizer (11). After centrifugation (15000 x g, 20 min, 4° C), supernatant was collected, and protein concentration was determined using BCA method (Bio-Rad, CA).

### Immunoblotting

Seventy microgram of whole brain whole cell lysate was run in a 1.5 mm thick 4%-12% Bis-Tris NuPAGE Novex system (ThermoFisher Scientific, MA) using NuPAGE MOPS SDS running buffer (ThermoFisher Scientific, MA) in the presence of 1:400 antioxidants (ThermoFisher Scientific, MA) first at 50 V for 30 min, then at 80 V for 30 min followed by 150 V for 1h on ice. Proteins were transferred onto a 0.2 μm PVDF membrane (Bio-Rad) at constant voltage 30 V for 1 h on ice using prechilled NuPAGE transfer buffer in XCeII II Blot Module (ThermoFisher Scientific, MA).

### Hippocampal slice preparation

Acute hippocampal slices (300 µm thick) were obtained from P16-P19 wild type and *Aif1*^*-/-*^ mice without differentiation of sex (12). All animals were anesthetized using 4% Isoflurane and sacrificed by decapitation. The hippocampi were isolated and cut using a VT1200s microslicer (Leica Microsystems Co.) in a solution containing (in mM): 215 sucrose, 2.5 KCl, 26 NaHCO_3_, 1.6 NaH_2_PO_4_, 1 CaCl_2_, 4 MgCl_2_, 4 MgSO_4_ and 20 glucose. Acute slices were placed in a chamber containing a 1:1 mix of sucrose cutting solution and extracellular artificial cerebrospinal fluid (A-CSF) recording solution containing (in mM): 124 NaCl, 2.5 KCl, 26 NaHCO_3_, 1 NaH_2_PO_4_, 2.5 CaCl_2_, 1.3 MgSO_4_ and 10 glucose at room temperature. After completing the slicing procedure, the 1:1 mix was replaced by A-CSF and slices were allowed to recover for at least 45 min prior to experimentation. For preparation of acute hippocampal slices from P90 mice, mice were perfused with 20 ml of cold NMDG solution containing in (mM): 93 NMDG, 2.5 KCl, 1.25 NaH_2_PO_4_, 30 NaHCO_3_, 20 HEPES, 25 glucose, 5 sodium ascorbate, 2 Thiourea, 3 sodium pyruvate, 10 MgCl_2_, 0.5 CaCl_2_, brought to pH 7.35 with HCl. The hippocampi were isolated and cut using a VT1200s microslicer in cold NMDG solution. Acute mouse slices were collected and placed in a chamber containing A-CSF solution incubated in a warm water bath 33-34°C. After 10 minutes of completing slice collection, the chamber was moved to room temperature and slices were allowed to recover for at least 45 min. All solutions were equilibrated with 95% O_2_ and 5% CO_2_ (pH 7.4).

### Electrophysiology

Whole-cell recordings were made from CA1 pyramidal cells voltage clamped at −65 mV using patch-type pipette electrodes (3-4 mΩ) containing intracellular solution (in mM): 131 cesium gluconate, 8 NaCl, 1 CaCl_2_, 10 EGTA, 10 glucose, 10 HEPES, pH 7.25 (280-285 mOsm). KOH was used to adjust pH. Series resistance (10-20 MΩ) was monitored throughout all experiments with a −5 mV, 80 ms voltage step, and cells that exhibited a series resistance change (>20%) were excluded from analysis. Miniature excitatory postsynaptic currents (mEPSCs) were recorded at 32 ±1° C in a submersion-type recording chamber perfused at 1-2 ml/min with ACSF supplemented with the GABA_A_ receptor antagonist picrotoxin (100 µM) and tetrodotoxin (0.5 µM) (13). Extracellular field excitatory postsynaptic potentials (fEPSPs) were recorded at 26 ±1° C in the presence of picrotoxin (100 µM) using a patch-type pipette filled with 1 M NaCl. A stimulating glass electrode was filled with ACSF and placed in *stratum radiatum* to activate Schaffer collateral inputs using a DS2A Isolated Voltage Stimulator (Digitimer Ltd.) with a 100 µs pulse width duration. Whole-cell voltage clamp and field recordings were registered with a MultiClamp 700B amplifier (Axon Instruments) and signals were filtered at 2 kHz and digitized at 5 kHz. Stimulation and acquisition were controlled with custom software (Igor Pro 6). Mini analysis (Synaptosoft) determined amplitude and frequency of mEPSC events. For extracellular field input-output experiments the DS2A isolated Voltage stimulator was increased from 0-25V, in 5V increments. Slopes of fEPSP responses and amplitudes of fiber volleys were determined using Igor Pro 6. Origin Pro 9 software was used to calculate slopes of linear-fit curves for input-output functions.

### Dendritic spine assessment

In whole-cell recordings of CA1 pyramidal cells, biocytin (0.2%) was included in the patch-pipette to load neurons for post-hoc morphological reconstructions (13). Acute slices from wild type and *Aif1*^*-/-*^ mice were fixed using 4% PFA in PBS 1X (pH 7.4) and further incubated with streptavidin-Alexafluor-647 for overnight at 4° C. Spine images were captured from both secondary and tertiary dendritic branches within 100 μm from soma by a laser scanning confocal microscope (Zeiss LSM880) assisted with the Airyscan module. Z-stack images were collected using a 63x objective at 0.36 μm interval for a total of 20 μm thickness and spine density assessed using the ImageJ software.

### Microglial morphology

Fourteen micron-thick fixed brain sections from P16-P19 wild type and *Aif1*^*-/-*^ mice were immunostained with anti-P2RY12 antibody and z-stack images acquired at 0.5 μm interval using a 60x objective with 2x magnification in an In vitro Ultima 2P microscope (Bruker Corp). Three hundred micron-thick fixed floating sections from adult wild type and *Aif1*^*-/-*^ mice were immunostained with anti-P2RY12 antibody and z-stack images acquired at 1.5 μm interval using a 25x objective in a laser scanning confocal microscope (Zeiss LSM880). Sholl analyses was performed by a researcher blinded to the conditions and genotypes by manually counting intersections through the concentric circles drawn incrementally at 5 μm intervals starting from soma (14, 15).

### Microglial motility in acute slices

Acute hippocampal slices from P16-19 wild type and *Aif1*^*-/-*^ mice were incubated with IB4-Alexa 594 (ThermoFisher Scientific, MA) for at least 30 min at a 25 µg/mL concentration as previously described (14). An In vitro Ultima 2P microscope (Bruker Corp) with an Insight Deep See laser tuned to 800 nm was used to visualize IB4-labeled microglia with 512×512 pixel resolution at least 50 µm from the surface in the *stratum radiatum* area of CA1 with 6-8 mW laser power measured at the 60x objective (Nikon, 1.0 NA). Identified microglia were imaged at 1X magnification generating a z-stack of 25 µm thickness with 1.7 µm steps to assess baseline microglia morphology. For motility induction a patch-type pipette containing ATP (1 mM) and Alexafluor-488 (30 µM) dissolved in A-CSF was placed in the *stratum radiatum* at least 50 µm from the surface. After acquiring baseline images of microglia at 800 nm, minimal pipette pressure was applied to puff ATP as previously described (14). Images were acquired at t=0 and at t=15 min. To analyze motility, Z-stack images of maximum intensity were compressed for baseline (0 min) and at 15 min using ImageJ software. The compressed images were merged and color coded-Red (0 min, baseline) and Green (15 min, after ATP application) using ImageJ software. To quantify directed motility, a circular region of interest (ROI) was drawn (100 µm diameter) around the pipette. Motility within the ROI was quantified to generate a directed motility index by using the RGB ImageJ plugin to determine the intensity of Red (0 min) and Green (15 min) pixels from merged images. The RGB values were used to calculate an index of motility ΔG=(G/R)-(R/G). When ΔG=0, a perfect alignment in pixels occurs indicating no net changes in pixel values, which suggested microglia experience nearly zero motility. When ΔG>0, pixels from G (t=15 min) outnumbered R (t=0 min) pixels in the field of view suggesting pixels were occupied by new microglia protrusions, new branch rearrangements, or retractions of protrusions seen at t=0 min (thus, R pixels are no longer occupied at t=15 min). When ΔG<0, pixels from R (t=0 min) outnumber G (t=15 min) pixels in the field of view suggesting there were few to no new microglia protrusions, branch rearrangements, or no retractions (thus, R pixels are maintained). Scorers determining directed motility quantifications were blind to genotype.

### Microglial CD68 quantification and synaptic engulfment

Thirty micron-thick fixed-frozen brain sections from wild type and *Aif1*^*-/-*^ mice were immunostained with anti-CD68, anti-PSD-95 and anti-P2RY12 antibodies (16). Z-stack images were captured at 0.33 μm interval, over a 7-8 μm depth, with 63x objective by a confocal laser scanning microscope (Zeiss LSM880) assisted with the Airyscan module. 3D-surface rendering and quantification of CD68 volume and engulfed materials were carried out blinded using the Imaris software. PSD-95 puncta only within CD68 structures were considered for engulfed materials which was performed by implementing the “ mask” function of Imaris as previously described (16, 17). To account for variations of cell size, the volume of CD68 as well as each engulfed material was normalized to each P2RY12 volume. Total PSD-95 volume per field from the same z-stacks was also calculated using similar protocol in Imaris.

### Behavior

All behavioral tests were performed using wild type and *Aif1*^*-/-*^ mice aged between 3-5 months. EthoVisionXT software (Noldus Information Technology Inc.) was used for tracking of mice for all behavioral assays unless otherwise mentioned. Experimenters were blinded to mice genotype information during data acquisition and analyses.

### Open field

Open field test is a general measure of exploratory/locomotor behavior as well as an index for anxiety was carried out as previously described (18, 19). Briefly, mice were allowed to move inside a box for 15 min during which their ambulatory movements were tracked. A zone defined as the center of the field which normal mice tend to avoid was assigned before the start of tracking.

### Elevated plus maze

The elevated plus maze test is a definitive test for anxiety-like behavior in rodents (20, 21). The test mouse was first placed at the center of the maze in a well-lit room, and its movement was tracked for 5 min. If a mouse jumped out, it was placed back into the maze. Data profiles include arm entries and risk management. Data were plotted as percentage time spent in the open arm over time spent in closed and open arm.

### Novel object recognition

Novel object recognition is a measure of cognitive ability in rodents (22, 23). Mice were first habituated by preexposure to an open field prior to the day of testing. On the test day, a mouse was first exposed to two identical objects by placing it at an equal distance from either object in the same open field and allowed to explore for 3 min. After an hour of delay (memory retention), the same mouse was re-exposed to the same open field for another 3 min now with one of the objects being switched to a new object. Exploration of old and new object was individually recorded using a software as well as manually. Percentage exploration for each object and total exploration time (for both objects) was calculated. The discrimination index was calculated by the formula: {T(Nov)-T(Fam)}/{T(Nov)+T(Fam)} during the time of testing.

### Social behavior

Social behavior assay was performed as previously described (16, 24). Mice were assayed in a Chamber (20 cm L x 40.5 cm W x 22 cm H, mouse sociability maze from Maze Engineers, Cambridge, USA) equally divided into three compartments. Dividing walls were constructed of clear plexiglass and at the center of each wall is a passageway that can be closed with a sliding door. In each of the outer compartments, there were two circular cages. Stranger mice belonged to same sex & approximately same age/weight, minimum 2 mice that were not housed in the same cage and in the same room as test mice. Stranger mice were habituated 2×5 min to the circular cage one day before testing. Test mice were transferred into cognitive room at least 30 min before testing for habituation (in home cages). There was a 5 min habituation trial for the test mice in the center chamber with closed doors after which they were transferred to the holding cage. A novel mouse was placed in the circular cage in the right adjacent chamber for all uneven testing mouse numbers while both doorways remained open. A test mouse was placed in the center chamber and allowed to explore all three chambers freely for 10 min after which the testing mouse was placed in the holding cage. A second unfamiliar mouse (not from the same home cage as the first stranger mouse) was placed in the cage in the remaining left chamber position (opposite first test) for all uneven testing mouse numbers (opposite for all uneven testing mouse numbers). The test mouse was placed back in the center chamber and allowed to explore all three chambers freely for 10 min after which all three mice were returned to their home cages. Sociability index was calculated by the following formula: {T(Stranger)-T(Empty)}/{T(Stranger)+T(Empty)}. The maze was cleaned with MB-10 to remove any trace scents from previous trials.

### Data analyses and Statistics

Data are presented as Average ± SEM. Statistical significance was determined by Student’s t test, or ANOVA with corrections for false discovery rate and with implementation of multiple comparison tests (such as Tukey’s or Sidak’s). Statistical analyses were performed in Graphpad Prism 8 and OriginPro 2015 (OriginLab). N represents total number of data points from at least 4 mice each genotype unless specified. Number of samples were balanced across genotype for a given set of experiments for appropriate statistical comparison. All analyses were performed by an experimenter blinded to the conditions and genotypes.

---

**Fig. S1.**
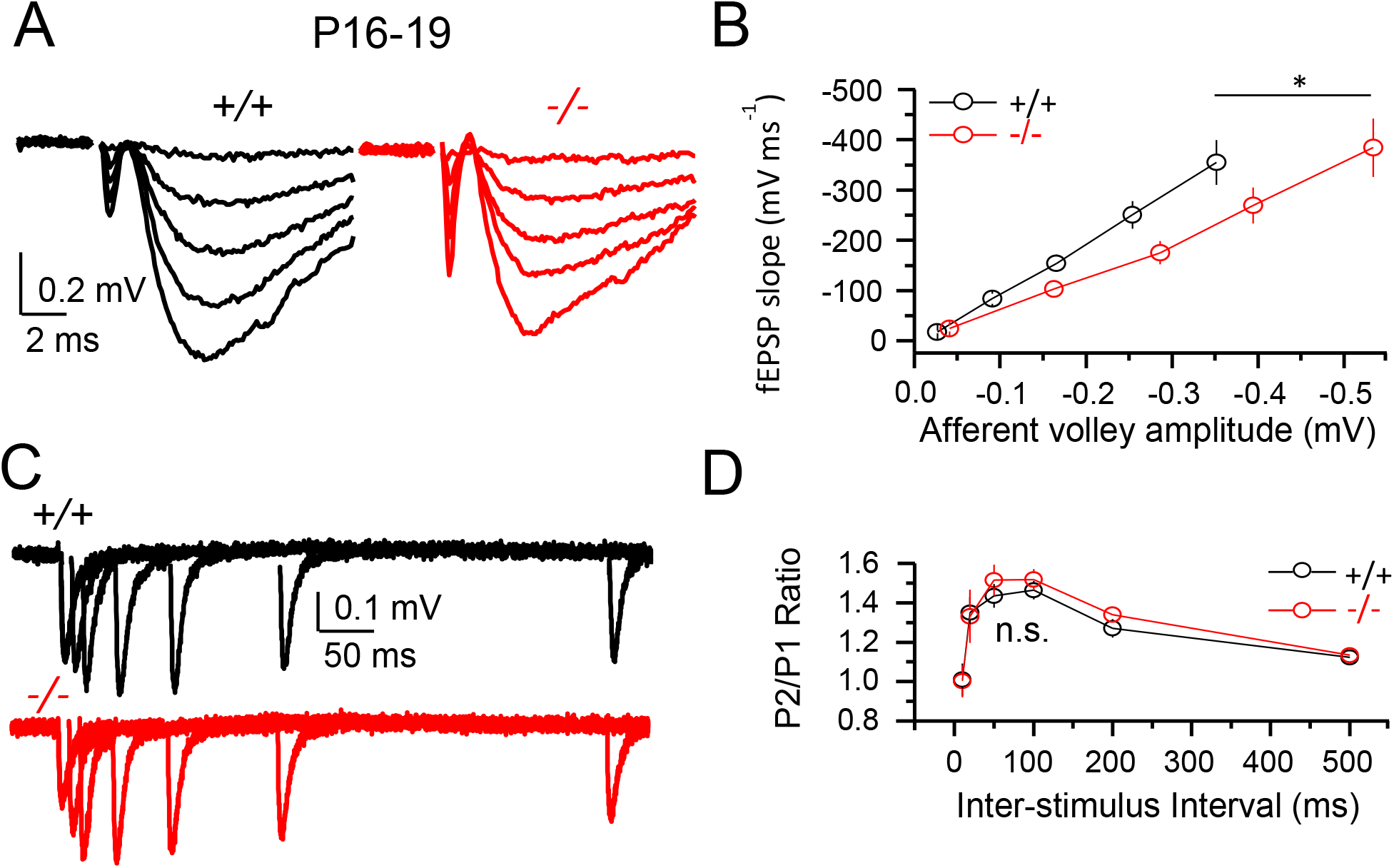
*Aif1*^*-/-*^ mice display reduced synaptic strength but normal paired-pulse ratio. *(A)* Input-output (I/O) curve of evoked excitatory postsynaptic potentials in CA3-CA1 synapses were generated from acute slices from P16-19 mice in the presence of picrotoxin (100 μM). *(B)* Slopes of fEPSPs plotted against fiber volley amplitude (Linear fit +*/*+: 1037.8 ± 13.59, r=0.99; *-/-*: 728.4 ± 34.85, r=0.99). n=10 slices from 4 mice per genotype; one-way ANOVA. *(C)* Superimposed traces from representative experiments for Paired-pulse ratio (PPR). *(D)* PPR was measured by dividing the slope of the second pulse by the first pulse (P_2_/P_1_). n=10 slices from 4 mice per genotype; one-way ANOVA. Average ± SEM. *, p<0.05. n.s.=not significant.

**Fig. S2.**
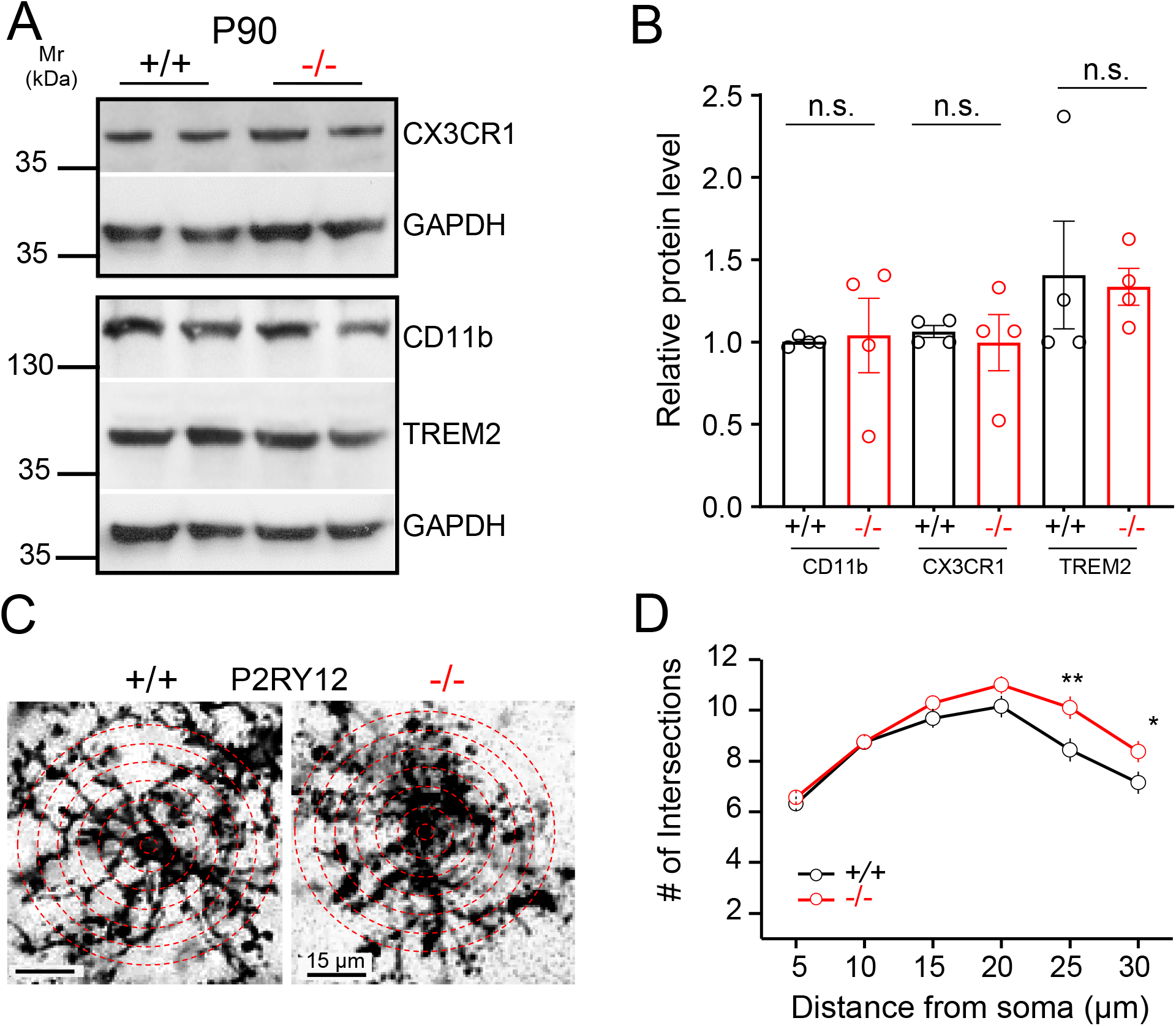
Normalization of postnatal microglial phenotype in the adult *Aif1*^*-/-*^ brain. *(A)* Western blotting of P90 whole brain whole cell lysates using antibodies against synaptic remodeling associated (CD11b, TREM2 and CX3CR1) proteins. *(B)* Quantitation of western blots in (*A*). n=4 mice per genotype, Unpaired Student’s t test. *(C)* Representative images of P90 fixed floating hippocampal slices immuno-stained with an anti-P2RY12 antibody. *(D)* Quantitation of microglial branch complexity revealed an over-compensation in P90 *Aif1*^*-/-*^ CA1 microglia. n=43-50 cells from 7 fields per region from 3 mice each genotype; Averages of every intersection were compared between wild type and mutant group in a two-way ANOVA correcting for multiple comparisons using Sidak’s multiple comparison test; F_1,539_=27.05; p<0.0001. Average ± SEM. n.s.=not significant.

**Fig. S3.**
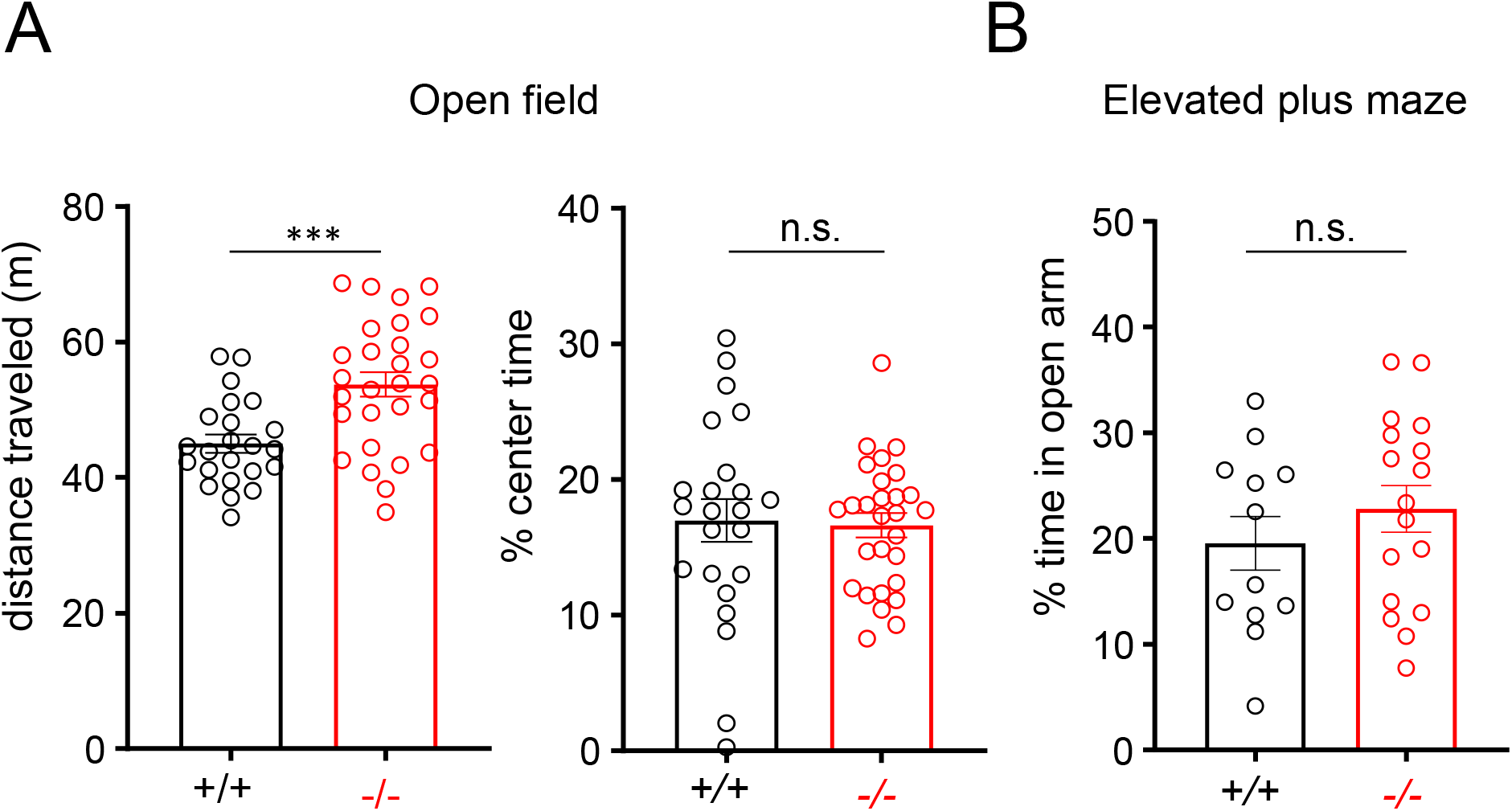
Adult *Aif1*^*-/-*^ mice exhibit enhanced exploration/locomotion but normal anxiety-like behavior. *(A)* Open field exploration was measured by the distance traveled (+/+: 45.0 ± 1.32 m; -/-: 53.74 ± 1.8 m; p=0.0004) and an anxiety-like behavior by the time spent in the center of the field during a 15 min trial. n=23 (+*/*+) and 28 (*-/-*), Unpaired Student’s t test. *(B)* Mice were allowed to explore an elevated plus maze for 5 min and time spent in open and closed arms was recorded. An anxiety-like behavior was assessed based on percentage time spent (+/+: 19.53 ± 2.53; -/-: 22.8 ± 2.2; p=0.34) in the open arm. n=12 (+*/*+) and 15 (*-/-*), Unpaired Student’s t test. Average ± SEM. ***, p< 0.001. n.s.=not significant.

**Fig. S4.**
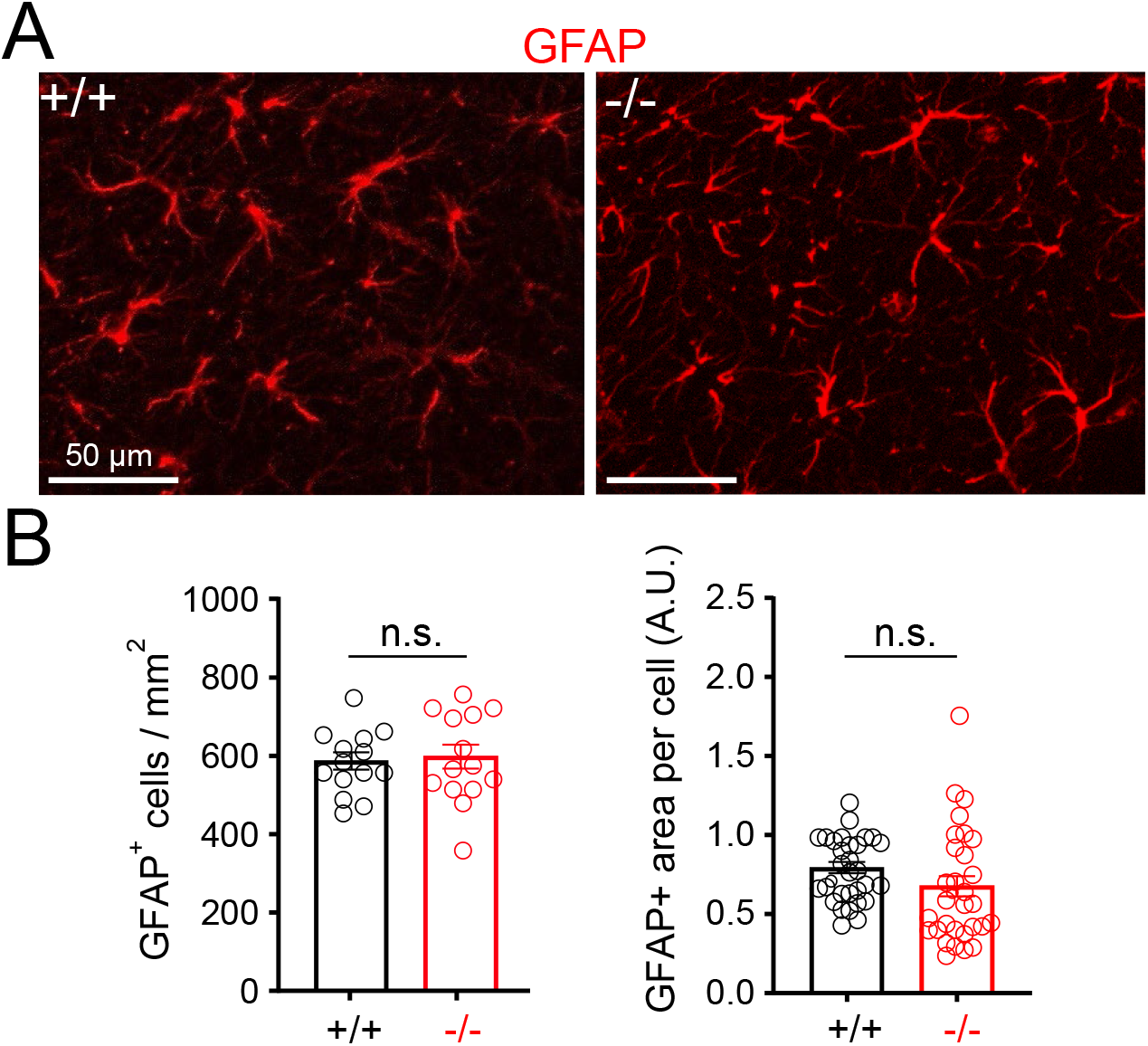
Normal astrocytic density and morphology in *Aif1*^*-/-*^ mice. *(A)* Representative confocal images of CA1 astrocytes immunolabeled with an anti-GFAP antibody in fixed frozen tissue sections from P16-19 mice. *(B)* Quantitation of astrocyte cell number (+/+: 568 ± 22 cells / mm^2^; -/-: 598 ± 31 cells / mm^2^; p=0.77) and area per cell reported by GFAP signal intensity (+/+: 0.79 ± 0.04; -/-: 0.68 ± 0.06; p=0.12) revealed no statistical differences between control and mutant groups. n=14 fields from 4 mice each genotype *(left)*; n=32 cells from 4 mice each genotype *(right)*, Unpaired Student’s t test. n.s.=not significant.

## Notes

### Competing Interest Statement

The authors have declared no competing interest.

